# Turgor and the conformational pathway for MscS recovery

**DOI:** 10.1101/2024.02.18.580915

**Authors:** Andriy Anishkin, Sergei Sukharev

## Abstract

The bacterial mechanosensitive channel MscS is an adaptive osmolyte release valve that cycles between closed, open, and inactivated states. Since some of these conformations are stable only in the lipid environment under specific conditions, the structures that are currently available cannot explain the entire functional cycle. Previous patch-clamp characterization has provided insights into the missing functional state by estimating protein expansion areas associated with the closed-to-open and closed-to-inactivated transitions and indicating that the closed state must be the most compact. In this paper, we model the conformational transition of MscS from the splayed conformation with the uncoupled gate to the putative compact closed state. The compaction pathway revealed in preliminary extrapolated motion simulations (ExMoS) involved an upward sliding motion of the internal TM3 barrel inside the outer sheath formed by TM1-TM2 helical pairs. This move leads to several structural changes: (1) the relocation of the characteristic kink at G113 to a new position at G121, (2) the establishment of the hydrophobic TM2-TM3 contact, (3) a new pattern of interactions with membrane lipids, and (4) the formation of stabilizing salt bridges between TM1-TM2 loops and the cytoplasmic cage domain. In the intact bacterial cell, the driving force for this upward motion is likely to be turgor pressure normal to the plane of the membrane acting on the upper hemisphere of the cage domain from the inside. Under continuing lipid synthesis in the inner leaflet of the plasma membrane, turgor pressure is also predicted to maximize the lateral pressure of lipids in the membrane, thus driving MscS compaction. Steered simulations were performed on the splayed state to mimic these effects by applying normal forces to the upper part of the cage domain and by applying lateral compression to the TM1-TM2 pairs, emulating the pressure of lipids. The structure arrived at the predicted compact state of the channel. This state was critically stabilized by displacing non-bilayer lipids from the TM2-TM3 crevices into the bilayer. We propose that the energized metabolic state of the cell generating high turgor and promoting lipid synthesis should strongly favor the compact closed state of MscS. The normal forces pressing the dome of the cage domain against the membrane may provide a common recovery mechanism for the entire family of MscS-like channels found exclusively in organisms with walled cells, which evolved to function under turgor pressure. A conversion of turgor into membrane tension under hypoosmotic cytoplasm swelling and peptidoglycan expansion will drive opposite processes of opening followed by adaptive MscS closure and inactivation.

## Introduction

The bacterial mechanosensitive channel MscS is the major osmolyte release valve initially identified in *E. coli* (Levina et al., 1999). It is ubiquitous in all free-living bacteria and is now considered critical for microbial survival in environments with changing osmolarity (Booth and Blount, 2012; Pivetti et al., 2003). The functions and the structural design of MscS turned out to be essential for all organisms relying on the presence of a cell wall. Thus, multiple members of the MscS family are found in representatives of all clades of walled organisms, including archaea, yeasts, protists, fungi, algae, and higher plants (Hamilton et al., 2015). Importantly, all these organisms rely on turgor pressure to maintain cell volume and shape. The rigidity and mechanical strength of the cell wall vary to satisfy the geometry of the pressurized cytoplasm and the magnitude of the hydrostatic pressure inside. Plant cell walls, typically withstanding 2-3 MPa (∼20-30 atm) of internal pressure, are probably the most rigid (Tyree and Hammel, 1972). Cell walls of gram-positive bacteria, yeast and fungi are also thick and strong enough to contain pressures of ∼15-20 atm (Atilgan et al., 2015; Money and Howard, 1996; Whatmore and Reed, 1990). In gram-negative bacteria, however, the single-molecular layer of peptidoglycan sandwiched between the inner and outer membranes is estimated to be more stretchable (ref), and the pressure inside an energized *E. coli* cell is estimated between 1.5 and 5 atm (Arnoldi et al., 2000; Cayley et al., 2000; Koch and Pinette, 1987).

Turgor pressure is hydrostatic pressure of an osmotic nature that is generated by the cell’s metabolic activity; it presses the membrane against the cell wall. Consequently, the cell wall experiences tensile stress, whereas the membrane normally feels only the normal compressive stress but no lateral tension. The obvious condition for ‘safe’ turgor pressure generation is the constant presence of some excess membrane area inside the cell wall that would exclude stretching under fluctuations of environmental conditions and cell growth. Since tensions exceeding 10-15 mN/m can damage the membrane, in addition to the pumps and synthetic processes generating turgor pressure, all walled organisms must possess mechanisms that would coordinate the growth of the cell wall with the replenishment of the cytoplasmic membrane material. The first support of the hypothesis of mechanical control of cell wall growth came from experiments with *Bacillus subtilis* (Rojas et al., 2017). In these microfluidic trials, the process of cell wall elongation was halted by stretching the cell osmotically, apparently by activating MS channels and temporarily de-energizing the cell.

The second indication for a mechanism intended to coordinate lipid synthesis with tension-sensing was the finding that the gene coding for the bacterial mechanosensitive channel MscM (*yjeP*) is in the same operon as the gene coding for phosphatidylserine decarboxylase (*psd*), the enzyme which completes the last step in PE synthesis. Both genes were found under the control of the periplasmic stress sigma factor σ^E^ and the Cpx two-component stress-signaling system (Hassoun et al., 2021).

Mechanisms controlling the integrity of the cell wall by adjusting turgor pressure to prevent damaging the cell wall and membrane must also be in place. Considering just gram-negative bacteria, one can imagine the situation when a bacterial cell wall is loosened after being subject to lysozyme or beta-lactam antibiotics (Wong et al., 2021). The less cross-linked and, therefore, more stretchable cell wall will give in and let the membrane completely unfold and experience tension. The mechanical stress in the system will be shifted from the cell wall resisting normal pressure to the cytoplasmic membrane, now acquiring lateral stress. In this case, the membrane itself becomes a stress-bearing element while reducing the normal turgor pressure. Notably, the lipid membranes are not designed to withstand substantial tension (ref). A similar situation occurs under osmotic downshock, where the osmotic influx of water increases turgor, stretches the elastic cell wall, and leads to the complete unfolding of the cytoplasmic membrane, thus generating tension. Safety mechanisms sensing both turgor pressure and the menacing conversion into membrane tension must exist. When tension builds up and opens mechanosensitive channels, MscS and MscL both act as safety valves; they release internal osmolytes, reduce osmotic gradient, and rescue cells from lysis (Booth and Blount, 2012; Cox et al., 2018; Kung et al., 2010).

Previously, we determined that MscS readily opens at super-threshold tensions, but under near-threshold tensions, it inactivates and stays inactivated as long as low tension persists (Akitake et al., 2005; Kamaraju et al., 2011; Moller et al., 2023). Now, we hypothesize that MscS is one of the devices capable of sensing both normal turgor pressure and lateral tension in the membrane, changing its state accordingly. Under turgor-dominating conditions, the channel must be ‘recharged’ to the closed ‘ready-to-open’ state. When the membrane unfolds completely and tension surges, MscS switches to the conductive ‘open’ state and fulfills its release valve function. When tension drops to a lower, near-threshold level, MscS transitions to the tension-insensitive and non-conductive inactivated state to completely reseal the membrane. When the cell recovers after the shock and regains its turgor, the MscS population returns to the closed state.

This sequence is based on functional patch-clamp studies of MscS that have revealed this adaptive functional cycle (Akitake et al., 2007; Boer et al., 2011). MscS inactivates from the closed state (Kamaraju et al., 2011). The separate closed-to-open and closed-to-inactivated transitions are both driven by tension, and each is characterized by its own kinetics, tension dependence, and a specific in-plane expansion, which present no puzzle from the point of view of mechanics or thermodynamics. However, the nature of the returning force driving the channel from the splayed inactivated state (with uncoupled tension transmission route) back to the compact closed state is still unclear.

The existing crystal and cryo-EM structures uniformly cluster around two conformations, the splayed non-conductive state (PDB IDs 2OAU, 7OO6) and the expanded state (PDB IDs 2VV5, 7ONL) with estimated partial conductance depending on the solubilization conditions (Flegler et al., 2021). Previous functional characterization of the closed-to-open and closed-to-inactivated transitions indicated that the closed state must be the most compact out of the three (Kamaraju et al., 2011).

While the tension-sensitivity of MscS has been well characterized previously (Belyy et al., 2010b; Britt et al., 2024; Sukharev, 2002), its dependence on normal turgor pressure has not been explored. To address how turgor pressure may influence MscS’s state distribution, we use computational approaches to study its putative conformational transition from the splayed inactivated state to the compact closed state. The initially used Extrapolated Motion simulations (ExMoS) suggested the upward sliding motion of the TM3 barrel inside the outer sheath of TM1-TM2. This motion relocates the characteristic kink in TM3 to a new position, establishes the predicted tension-transmitting hydrophobic TM2-TM3 contact (“clutch”) (Belyy et al., 2010a), and re-forms stabilizing salt bridges between TM1-TM2 loops and the cytoplasmic cage domain. Steered simulations performed under normal forces mimicking turgor and simultaneous lateral compression of peripheral helices mimicking the pressure of lipids resulted in a very similar compact state of the channel as generated by ExMoS. We predict that the energized metabolic state of the cell generating high turgor should favor the return of MscS to the closed state. This may be a common recovery mechanism for the entire family of MscS-like channels.

## Methods

### Extrapolated Motion simulations (ExMoS)

The ExMoS was designed (Anishkin et al., 2008a; Anishkin et al., 2008b) to explore the sterically permitted directions of protein domain motions quickly. The simulations are implemented as scripts in VMD using bare protein structures in a vacuum (with charges adjusted to reflect exposure to solvent). The procedure illustrated in Fig. S1 starts with an initial displacement (0.1-1 A) of all atoms in an entire protein or a specific domain in an arbitrary direction or just as a random thermal fluctuation. The first energy minimization corrects the inter-atomic distances and removes possible steric clashes; the following short MD simulation additionally relaxes the system by overcoming possible entrapment in local minima. After the simulation, the originally symmetric conformation of a multimer may become slightly asymmetric due to the application of random forces in the Langevin temperature control algorithm. If the pathway is desired to be symmetric (which usually improves the stability of the system and decreases thermal defects), an optional symmetrization step can be included afterward, and symmetry restraints can be periodically imposed during the simulation. The final energy minimization completes the iteration to produce the current set of coordinates. The next cycle begins with the displacement of the atoms implemented as the linear extrapolation of coordinates based on the previous and current values. Once displaced, energy minimization, relaxing MD simulation, symmetrization, and the second minimization are repeated.

**Fig. S1.**
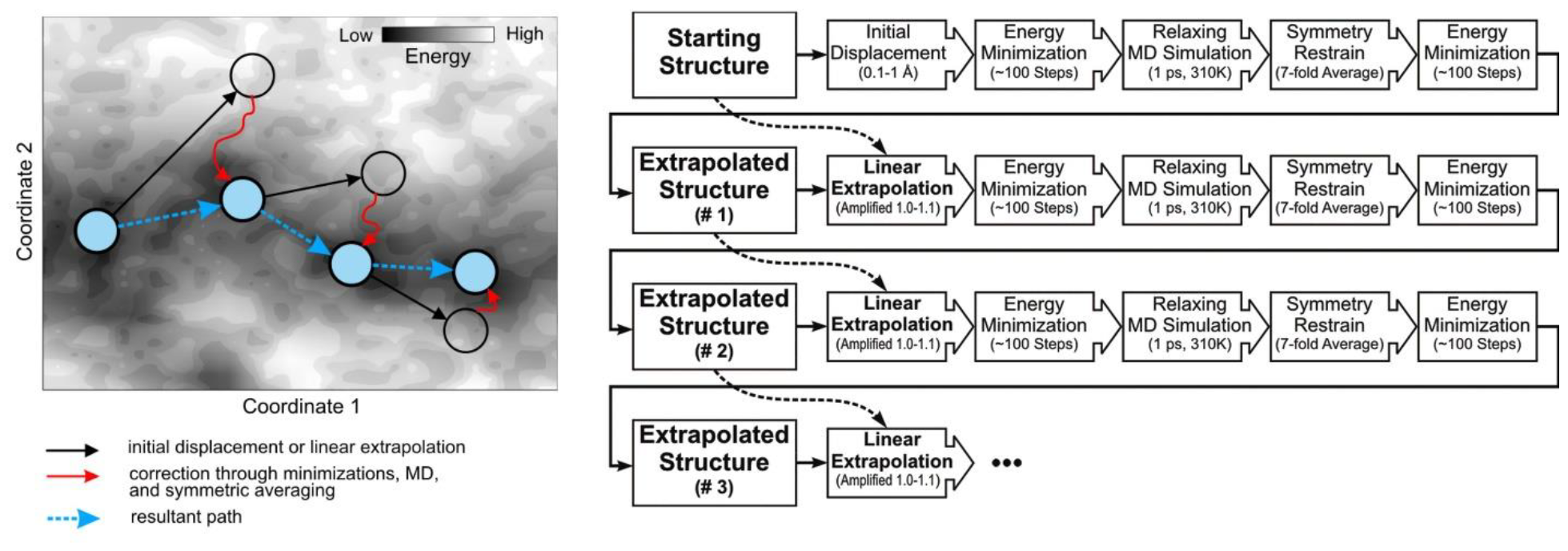
The principle of the procedure and extrapolation-minimization-relaxation cycles constituting the extrapolated motion protocol.

In our extrapolated simulations specifically for MscS, random starting 0.1-1 Å displacements due to thermal fluctuations were used to initiate expansion or compression. In each cycle, the extrapolated displacements of atoms were ‘corrected’ in their absolute value and direction by the initial energy minimization (100 steps, conjugate gradient, and line search algorithm), a 1 ps relaxing MD simulation at 310 K and completed by another energy minimization (100 steps) with seven-fold symmetry restraints. To speed up or slow down the extrapolated motion, the magnitude of extrapolated displacement at each step could be adjusted by introducing a linear ‘amplification coefficient’ (γ), increasing the absolute value of the displacement vector. It was determined empirically in multiple trials that at γ = 1 (series of un-amplified displacements), the initiated motion of the barrel dampened quickly, the expansion slowed down and transformed into low-scale (1-2 Å) oscillations without reaching the diameter expected for the open state. At γ > 1.2, the structure often expanded beyond the point of instability. A resultant set of discrete states could be approximated with a smooth quasi-continuous trajectory covering a meaningful range of the transition. When merged into a continuous trajectory, ∼50 extrapolation cycles typically produced a substantial displacement. Because extrapolations have a random component due to the Langevin temperature control, when repeated from the same starting conformation, independent series were observed to produce varying trajectories, usually with similar features. Performed in vacuum, the extrapolated transitions yielded models that have to be further tested and refined in all-atom MD simulations (below).

### Equilibrium simulations with symmetry-driven annealings and the statistics of contacts

The resting state best satisfying the experimental parameters was MD simulated in a fully hydrated POPE-POPG (3:1) bilayer (220 lipids) with TIP3P water (5) as NPT ensemble at 310 K (Langevin method), in a flexible hexagonal periodic box at 1 atm pressure (Langevin piston method), PME electrostatics, 12 Å cutoff for VDW and direct electrostatic interactions, and a membrane tension of 10 dyne/cm.

Potassium and chloride atoms were added in amounts corresponding to 200 mM salt concentration. The total system size was about 168,000 atoms. After 150 ns unrestrained simulations, they were followed by 1 ns annealing to clarify the overall trends in conformational change. During the annealing stages, all the protein atoms were gradually driven towards the continuously updated 7-fold symmetric average positions using harmonic restraints with spring constant increasing exponentially from 0.001 to 20 kcal/mol/Å2. The rest of the system (lipids, water, and ions) was unrestrained.

### Steered MD

Steered MD simulations were performed in an all-atom system to explore whether externally applied forces imitating the effect of turgor on MscS in the inactivated conformation will cause conformational changes similar to those observed in the ExMoS transition to the resting state model. The starting system was taken at the symmetry-annealed conformation after a 150 ns unrestrained simulation started from 6pwp as initial conformation. All the simulation conditions and settings were the same as in the equilibrium simulations.

We explored the effect of two kinds of steering forces: 1) the upward pressure on the central region of the top of the cage (center of mass of the backbone of residues 105 to 109) and 2) the lateral compression of the cytoplasmic part of TM1-TM2 helical couple (residues 35 to 87 harmonically steered towards the pore axis). To prevent the spatial drift of the protein under the action of the steering and thermal forces, the backbone of the periplasmic residues 13 to 31 was harmonically restrained near the starting positions. We explored the effects of two steering forces: 1) the upward pressure on the central region of the top of the cage (center of mass of the backbone of residues 105 to 109) mimicking turgor pressure was simulated for 30 ns to achieve steered motion and displace the gate 15 Å upward, and 2) the lateral compression of the cytoplasmic part of the TM1-TM2 helical pair (residues 35 to 87 harmonically steered towards the pore axis) representing the lateral pressure of lipids was simulated for 12 ns under a harmonic constant ramp. The two motions were performed either separately (only gate displacement or only compaction) or in sequence as follows: first, an upward displacement of the gate, then an optional step of repositioning of the wedged lipids from the crevices between TM1-TM2 and TM3 helices, and then the lateral compaction of TM1-TM2 helices. We also tried simultaneous combinations of gate displacement, optional lipid removal, and TM1-TM2 compaction. Each combination of simulations, upon completion, was followed by a 1 ns symmetry annealing simulation to reveal the dominating structural trend. To prevent the spatial drift of the protein under the action of the steering and thermal forces, the backbone of the periplasmic residues 13 to 31 was harmonically restrained near the starting positions.

## Results

### 1. Extrapolated Motion Simulations (ExMoS) suggest the compaction pathway

Extrapolated motion simulations were performed using the same protocol as described in our earlier publications on the modeled structures of the closed (Anishkin et al., 2008a) and open (Anishkin et al., 2008b) states, with the difference here that these systems were initiated from a slightly different inactivated conformation (6pwp) with the N-terminus modeled based on our earlier representations. The reason for re-modeling the N-terminus rather than using it as resolved in the 6pwp structure was that in unrestrained all-atom simulations, the N-terminus quickly became disordered and lost contact with the membrane, whereas it had a much more stable fold in our earlier models of the inactivated, closed, and open states (Anishkin et al., 2008a; Anishkin et al., 2008b). While trajectories in the current set that were started from the splayed, inactivated conformation often tended to expand the transmembrane barrel, some of them instead showed compaction such that TM1-TM2 helical pairs approached the central TM3 barrel and led to its displacement (including the gate region) towards the periplasmic side, usually by a distance comparable to a couple of helical turns. The upward motion also resulted in the re-formation of one more helical turn in the upper part of TM3 (residues 92-94), which is uncoiled in all splayed conformations. The overall compact state was similar to the previously published resting state model (Akitake et al., 2007; Anishkin et al., 2008a), including such features as the straightening of the helical kink around G113 and the formation of the kink near G121, which essentially allowed the upward displacement of the gate along the pore axis and establishment of hydrophobic contact between TM1-TM2 pairs and TM3s.

### 2. Stability of the compact state in equilibrium MD simulations

Both structures, 6pwp and the compact model, were equilibrated in the POPE/POPG bilayer for 10 ns with a restrained backbone, then relaxed for 10 ns without restraints, and then simulated under 20 mN/m for 40 ns. The drift of the 6pwp structure in the transmembrane region (backbone of the residues 21 to 127) was 4.03 (RMSD), whereas the transmembrane domain of the compact model slightly relaxed in the outward direction (2.93 RMSD). The relatively rigid cage domain experienced no significant distortions in either case.

The compaction procedure brought R59 closer to D67 on the adjacent subunit, and the TM1-TM2 connecting loops approached the R128 and R131 pairs of arginines located on the upper hemisphere of the cage. The more splayed conformations prefer the D62-R128 salt bridge, whereas the more compact conformation favors the D62-R131 salt bridge. As a result, the compaction produced a network of fourteen peripheral salt bridges, stabilizing the tightly folded state with more upright peripheral helices. The dynamics of these salt bridges in the simulated compact state are shown in Fig. 2. As the higher RMSD of the splayed structure suggests, TM1-TM2 pairs are noticeably more dynamic than in the compact model. This leads to a lower overall number of salt bridges in the splayed structure. The 6pwp structure strongly preferred the D62-R128 salt bridge, as shown in panel B. The number of this type of bridge fluctuated only between 1 and 2 out of the whole heptamer. In contrast, the more stable compact model had the total number of bridges fluctuating between 6 and 9, with the dominant contribution from the D62-R131 pair. On average, each subunit had at least one of the bridges, thus stabilizing the whole TM domain packing.

**Fig. 1.**
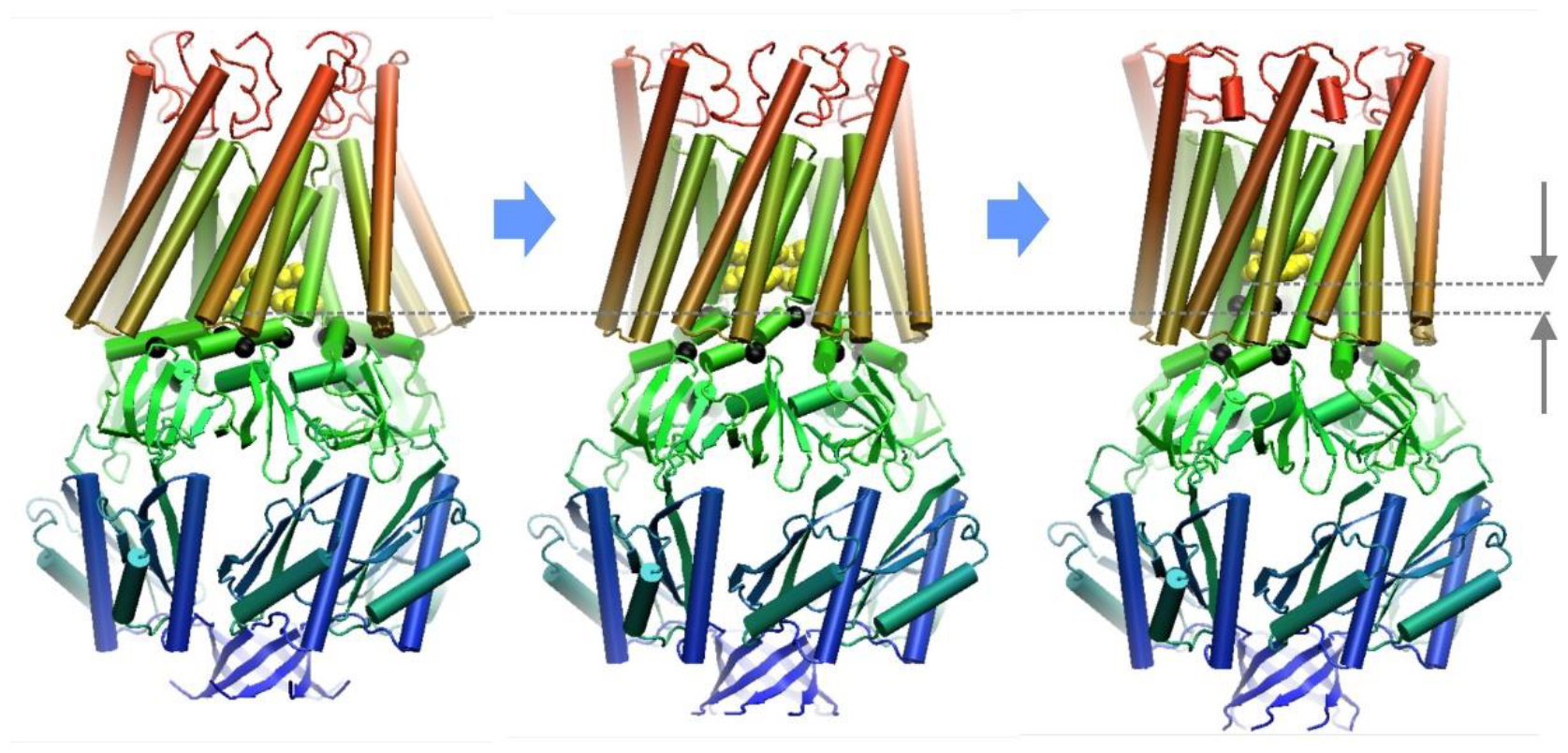
Extrapolated motion simulations (ExMoS) of the original 2OAU structure with modeled N-termini. The snapshots represent a typical extrapolated trajectory of compaction with the starting, intermediate, and end conformation, resulting in a putative closed state with established TM2-TM3 contacts, an upward-shifted gate (arrows on the right), and new positions of TM3 kinks. Hinge glycines, G113 and G121, are shown as black beads.

**Fig. 2.**
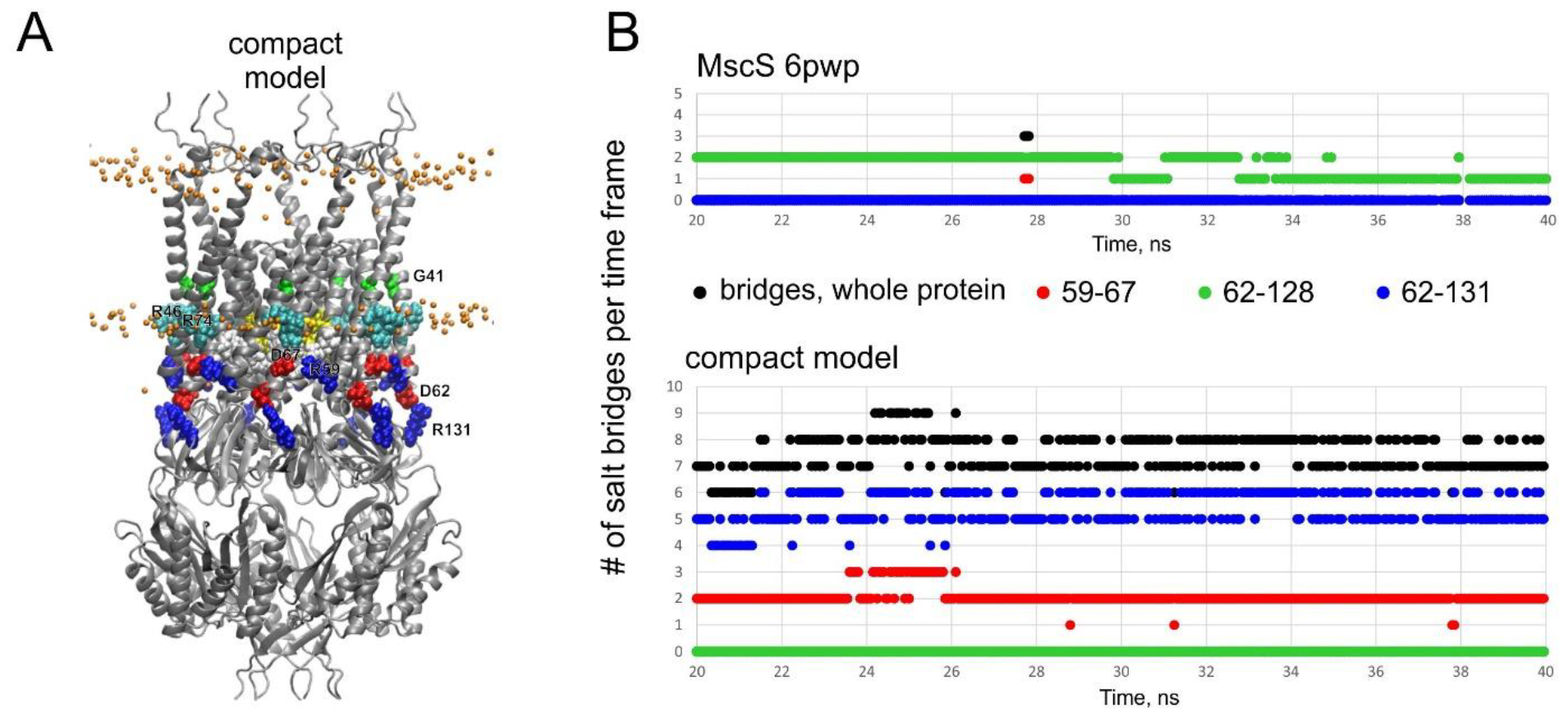
The structural properties and the stability of the compact state. (A) Established peripheral salt bridges R59-D67 and D62-R131. (B) Probabilities of peripheral salt bridges in the simulated 6pwp structure and in the compact closed state model.

The clear difference between the splayed and compact conformations is the accessibility of the pore-forming TM3 to the membrane lipids. All available non-conductive structures have a kink at G113, dividing the TM3 helix into the TM3a and TM3b segments. In several structures (Flegler et al., 2021), and simulations (Pliotas et al., 2015), lipids intercalating between the TM-TM2 pairs are pulled out of the membrane and acquire a non-bilayer orientation. Additional lipids are frequently found positioning their headgroup near R88 at the TM2-TM3 turn facing the periplasm (Flegler et al., 2021; Reddy et al., 2019). Lipid contacts with TM3b segments were scored throughout the simulation and plotted in Fig. 3B. The plot shows the probability of a contact of a particular sidechain with any of the lipid atoms. As one can see, the 6pwp structure shows contacts all the way up to the end of TM3b. The contacts with TM3b are due to the presence of non-bilayer lipids in the pockets located already below the lipid boundary. Remarkably, the compact structure completely excludes lipid contacts with the cytoplasmic side of TM3, allowing for contacts primarily above L111.

**Fig. 3.**
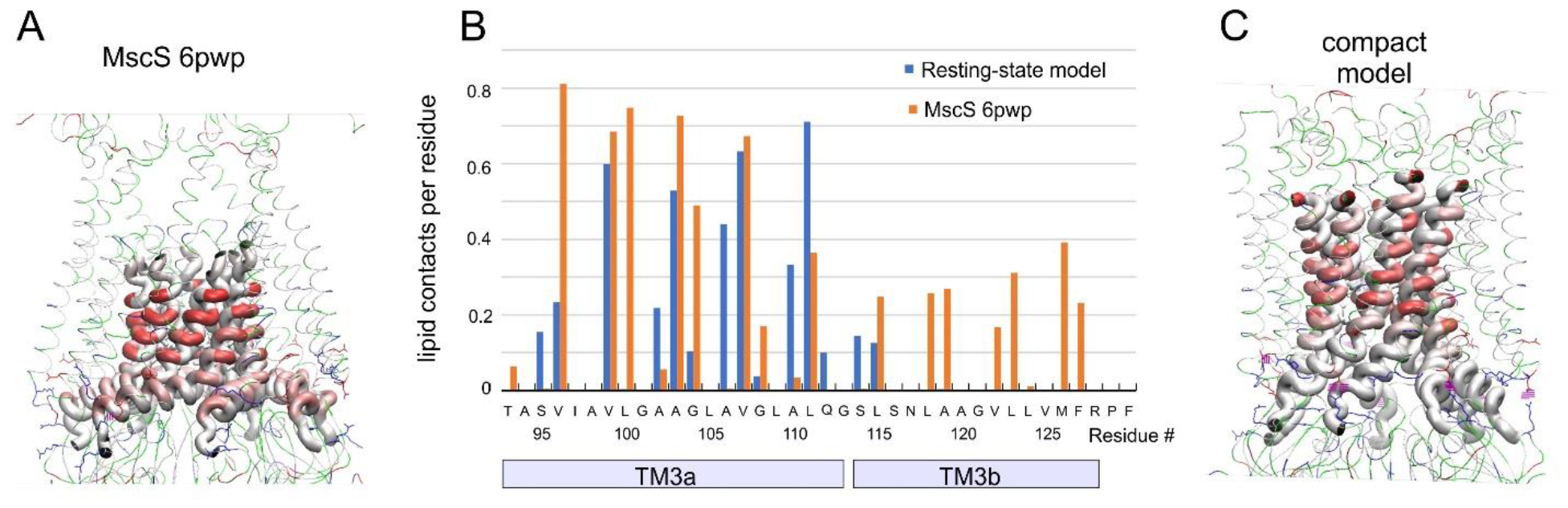
Equilibrium MD simulations of the compact structure. Lipid accessibilities of the TM2 and TM3 helices (B) in the splayed (6pwp) (A) and compact conformations (C).

### 3. Analysis of mechanical forces that could drive compaction

The hollow cytoplasmic domain (cage) has seven portals connecting its interior with the cytoplasm. The cage is rigid, and the portals have elongated shapes about 7×14 angstroms in size, allowing it to pass water, inorganic ions, small compatible osmolytes (glycine betaine), organic anions (glutamate, succinate), mono-sugars, and trehalose. Larger osmolytes and macromolecules are excluded from the cage and create ‘crowding’ pressure acting from the outside. Previous experimental and computational analyses have shown that external crowding pressure drives fast inactivation under moderate membrane tension. Turgor pressure transmitted through the solvent (aqueous solution), in contrast, acts uniformly from outside and inside the cage. As illustrated by simulations of the lipid-embedded structure (6pwp), water is everywhere; it is in the cage and has a large normal component pressing the upper hemisphere into the membrane. Importantly, water is not present in the area outside the cage covered by the transmembrane helices. Therefore, the dominating pressure will act on the upper hemisphere in the upward direction. Additionally, water pressure acting from the sides will close the gaps between TM2 and TM2. Turgor appears to be one of the important forces that work in the direction of the predicted barrel motion.

### 3. Steered simulations of splayed MscS with localized forces mimicking turgor and lateral lipid pressure produce the compact state

Given the trends in ExMoS towards spontaneous compaction of the TM1-TM2 domains around the TM3 barrel and the movement of the gate-bearing TM3a towards the periplasmic side of the channel, we tested whether the expected turgor-related forces applied to the top of the cage domain in the vicinity of the gate region (the backbone of the residues 105 to 109), and/or lateral compaction of TM1-TM2 paddles at the cytoplasmic side of the channel (backbone of the residues 35 to 87 as a reflection of combined lateral water and lipid pressure) would cause similar conformational changes in an all-atom system with explicit solvent and membrane. As described in more detail in the Methods section, we have explored these two steering strategies both separately and in sequential combination. When applied separately, these steered strategies did not lead to a conformation similar to the ExMoS-generated model of the compact resting state. 30 ns gradual displacement of the harmonically retrained gate region towards the periplasmic side by 15 Å did produce a gate position similar to the modeled conformation. However, it did not lead to the compaction of TM1-TM2 around TM3s and did not cause the exit of lipids from the crevices. Moreover, during the following 1 ns symmetry annealing, the structure quickly returned to the starting conformation resembling 6pwp. Gradual 12 ns lateral compression of the cytoplasmic part of TM1-TM2 paddles led to a much more compact structure. However, it did not cause any displacement of the TM3a barrel towards the periplasmic side, nor did it expel the lipids from the crevices. The latter might not be surprising because, in experiments, the recovery from inactivation (which, according to our model, requires the removal of the lipids from the crevices to establish direct contact of TM1-TM2 with TM3) occurs on the timescale of seconds, which is 8 orders of magnitude longer than the times we used to explore the conformational space in the current steered simulations. The most effective steering strategy turned out to be a two-step transformation: (1) upward motion of the gate, (2) removal of lipids from the crevice, and (3) application of compressive forces to the TM1-TM2 paddles. It repositioned the gate to the intended position, straightened the kink at G113, established a kink in the new position at G121, and formed a contact between TM1-TM2 paddles and (now elongated) TM3 barrel. Importantly, the presence or absence of lipids in the crevice was a crucial factor for the stability of the assembly. When lipids were still present (although partially expelled from the crevices during the lateral compaction), the structure quickly returned to the splayed conformation. However, if the lipids in the crevices were manually removed and repositioned into the bulk after the upward steering of the gate and before the compaction, the final compacted structure had a smaller lateral footprint, tighter contact between TM1-TM2 pairs and TM3 barrel, and was very stable during the symmetry annealing and the following unrestrained equilibrium simulations testing its stability.

Previous studies proposed that tension pulls out the lipids from these cavities, and this results in channel opening (refs). The comparison of volumes taken by the protein-lipid system in the splayed and compact states and the experimental observation that membrane stretch impacts recovery from inactivation dictates that pressure should squeeze the lipids out of the crevices and put them in a more compact (lower footprint) conformation with their peers in the bulk of the bilayer. Lateral compressibility of lipids in the bilayer is predicted to be higher in the bilayer than in between the helices. The more expanded cytoplasmic side of the transmembrane domain will be more sensitive to lateral pressure in the inner leaflet.

## Discussion

The adaptive functional cycle of MscS includes at least three states: closed, open, and inactivated. The separate tension-dependent opening and inactivation transitions both originate from the closed state. From experimental estimations (Kamaraju et al., 2011), the closed state has a projection area to the plane of the membrane that is about 9 nm^2^ smaller than the inactivated state and, therefore, is predicted to be the most compact. The existing structures do not explain the functional cycle. They uniformly cluster around two conformations, the splayed non-conductive and the presumably conductive expanded state, with the estimated conductance that accounts for 50-60% of the fully open state. The two states are easily interconvertible based simply on the solubilizing conditions (Flegler et al., 2021). The partially conductive expanded state does not exist in regular lipids, only in detergents (Lai et al., 2013) or short-chain lipid nanodiscs (Park et al., 2023; Zhang et al., 2021).The absence of structures with physically meaningful parameters of the in-pane area, conductance, and interhelical connectivity prompted our modeling effort.

The previously performed ExMoS predicted an upward sliding motion of the TM3 barrel in the outward direction. This movement is apparently limited by two structural features: the length of the periplasmic TM2-TM3 loop, which partially unfolds when the barrel moves down, and the position of the TM3 kink on the cytoplasmic side that limits the upward motion. The latter obstacle is overcome by straightening TM3s at G113 and moving the kink to G121, which is the most conserved residue in the entire family of MscS-like channels. The result is the more upright conformation of the TM1-TM2 pairs, restored hydrophobic contact, and re-formed salt bridges. We propose that intracellular turgor pressure acting on the upper hemisphere of the cage and the gate region is the main driving force that moves the TM3 barrel.

The physical forces that can drive this type of transition include turgor pressure and lateral pressure of lipids. Turgor pressure directed normally to the plane of the membrane is predicted to assist in the outward motion of the TM3 barrel. Simultaneously, lateral pressure of lipids should re-orient the splayed TM1-TM2 pairs and move them closer to TM3s. In essence, the lateral compression of lipids is also the result of the turgor pressure acting on the entire cytoplasmic membrane, including the vicinity of the channel, that pushes it against a more rigid cell wall. The natural situation that embraces this scenario is the osmotic shock and recovery cycle illustrated in Fig. 7.

The results of the simulations presented above also emphasize the dual role of the cage domain. Previously, we have shown that increased crowding pressure acting on the outer surface of the cage drives fast inactivation when MscS is subjected to moderate near-threshold tension (Rowe et al., 2014). Large, portal-impermeable cytoplasmic components exert this crowding pressure. We should note that the osmotic permeability response to hypotonic shock leads to a massive release of osmolytes followed by water efflux and thickening of the cytoplasm. This crowding-driven inactivation protects the cell from excessive dehydration by stopping the efflux of osmolytes and water. When the permeability response is over, the cell regains its metabolic activity, rehydrates, and restores its volume and turgor.

MscS’s response to turgor again engages the cage but in a different way. In this case, it is not the outer surface of the cage facing the cytoplasm that is involved, but rather the inner surface closest to the gate. Water, dissolved ions, and small organic osmolytes easily equilibrate between the inside and outside of the cage and exert hydrostatic pressure in all directions.

Importantly, water conveying this hydrostatic pressure throughout the cytoplasm and cage is completely absent in the lipid-filled TM2-TM3 crevices in the splayed state (Fig. 4). Thus, the compacting action of turgor pressing the dome from the inside of the cage will be unopposed. With this regard, the pressure-receiving dome adjacent to the gate region would function as a ‘piston’ driving the TM3 barrel inside the firmly lipid-anchored TM1-TM2 sheath (Britt et al., 2024).

**Fig. 4.**
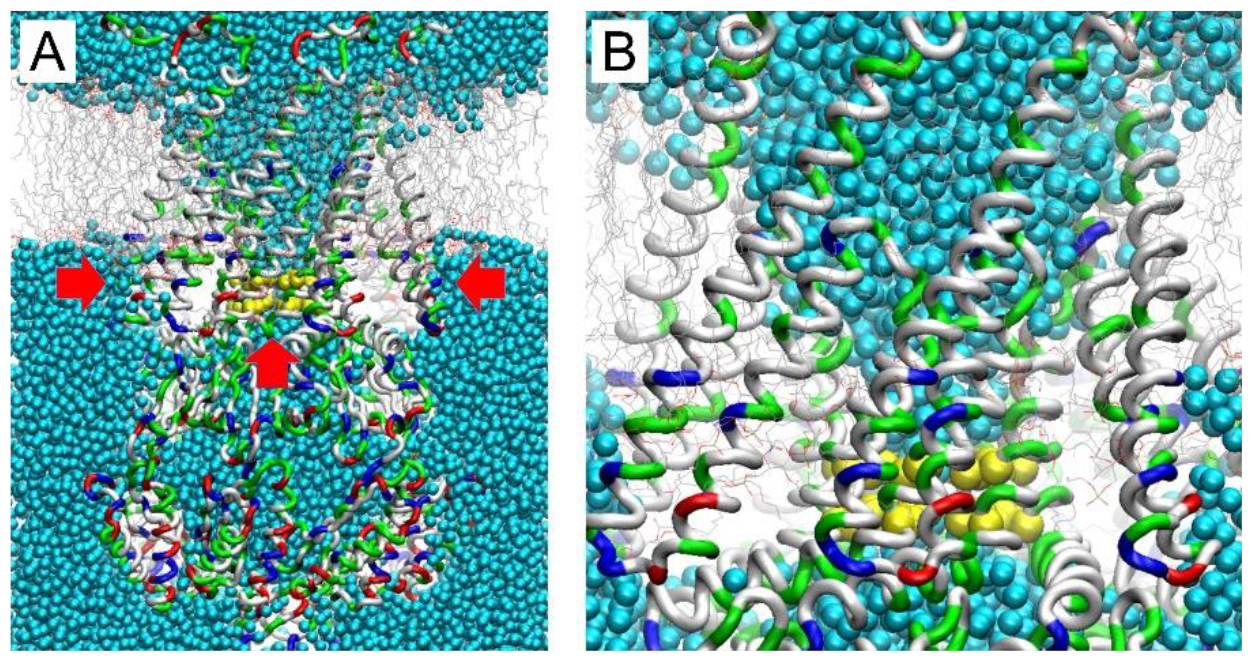
Equilibrium MD simulations of the splayed 6pwp structure reveal water present outside, everywhere inside the cage, and largely solvate the pore. Importantly, TM2-TM3 interhelical crevices are completely desolvated. The picture shows a 20 A vertical slab across the structure at the end of the 40 ns simulation. (A) Full view, (B) zoom on the crevices. The direction of pressure acting on the gate region from all sides is shown by red arrows.

**Fig. 5.**
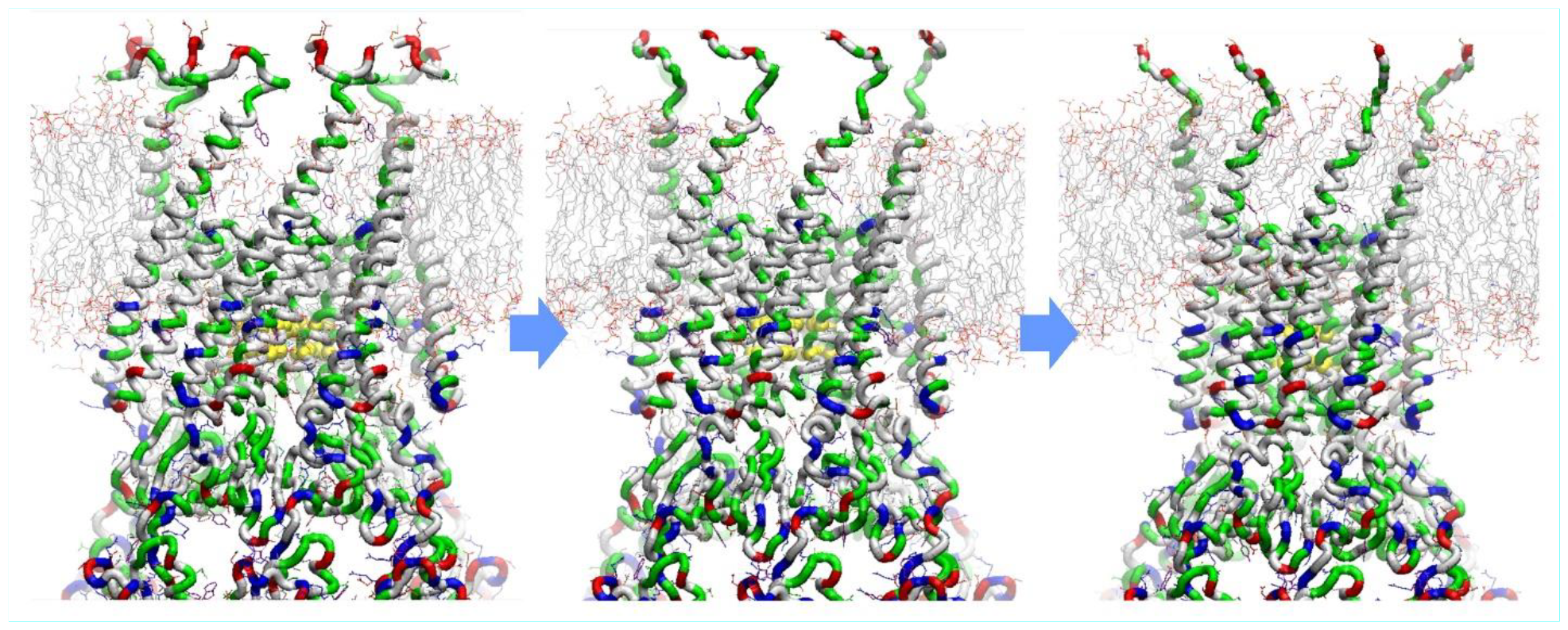
Illustration of steered simulations with a three-step compaction procedure: (1) upward movement of the gate, (2) removal of lipids from the crevices to the bulk of the bilayer, and (3) compression of the cytoplasmic ends of the concomitant removal of lipids from the crevices . After the first stage, the removal of lipids from the crevices dramatically stabilized the final structure.

**Fig. 7.**
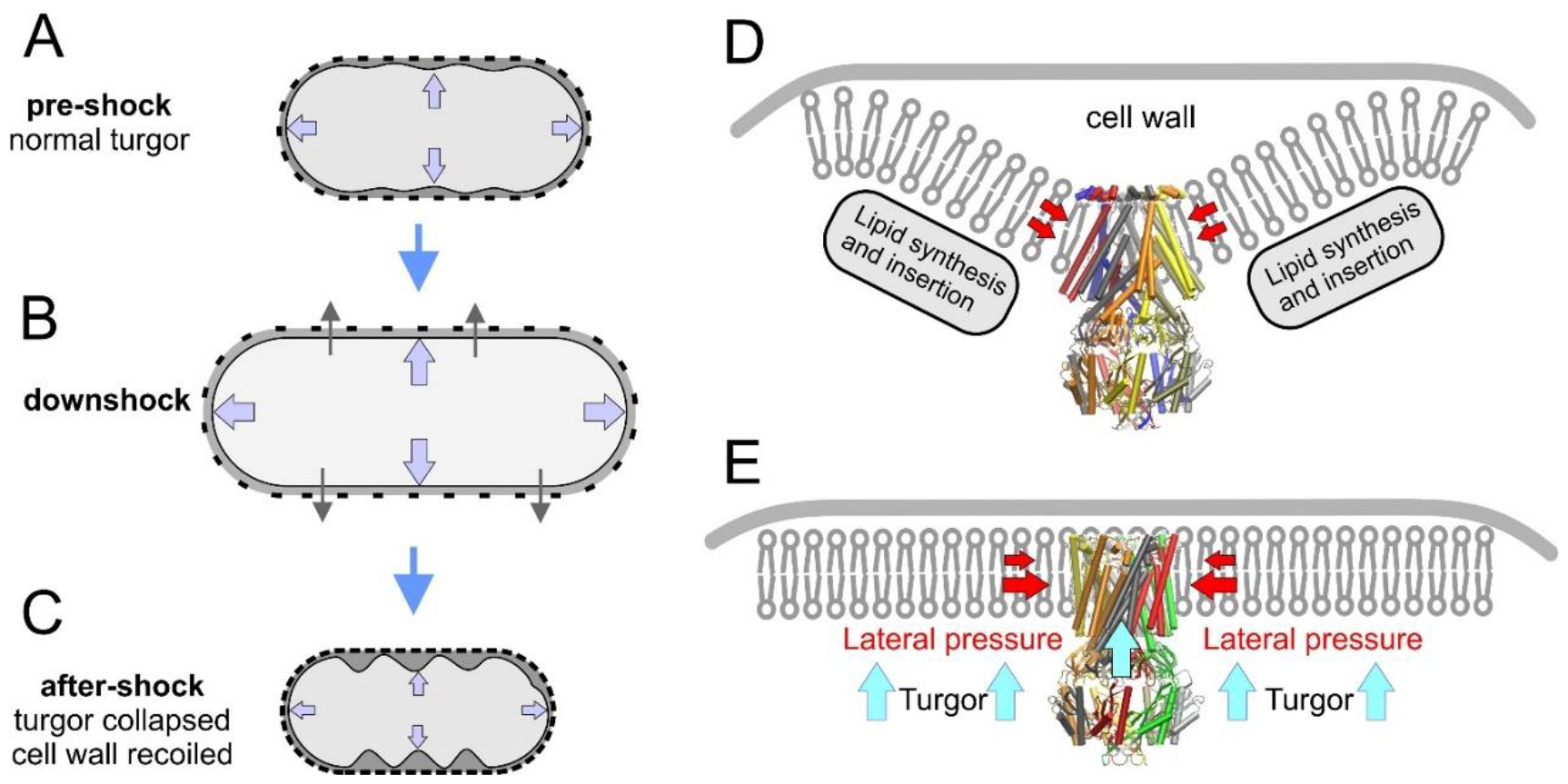
The mechanical state of a bacterial cell during shock-recovery events and the related distribution of MscS population between different functional states. (A-C) The events occurring to a bacterial cell during the hypotonic shock are a normal turgid state before the shock (A), hypotonic swelling upon media dilution and osmolyte release phase (B), cell deflation, and recovery (C). The picture zoomed on a single MscS channel, and its immediate environment inactivated post-shock state in a wrinkled membrane (D) and a recovered state after regaining turgor by the cell (E).

A constant interplay exists between metabolically generated turgor pressure, peptidoglycan elasticity, and membrane area inside the cell. Turgor does not necessarily create tension. If the membrane inside the peptidoglycan layer has an excess area, it will simply be pressed against the sacculus. Tension will be generated under osmotic downshock when excessive turgor stretches the peptidoglycan elastically, and the membrane is completely unfolded (all extra area is used up). A similar effect is expected when the cell wall is compromised (nicked or partially digested). It becomes more stretchable in this case, and full cytoplasm swelling will be observed even at normal turgor pressures. Complete membrane unfolding generates membrane tension followed by channel activation and partial osmolyte release. If moderate tension persists, MscS will inactivate. Inactivation is a defensive action to prevent intermittent channel opening and to reseal the membrane under non-lytic tensions. Therefore, the elasticity of the peptidoglycan determines the point at which turgor pressure (normal to the cell wall) is converted into lateral tension in the cytoplasmic membrane. At this moment, MscS, having lost the normal force that maintains the closed state, now receives the lateral stimulus that opens and inactivates the channel. This illustrates that the popular ‘force-from-lipids’ concept (Martinac and Kung, 2022) does not provide a complete set of parameters describing MscS gating, and bulk forces must be considered.

We conclude that the energized metabolic state of the cell generating high turgor (and possibly hyperpolarization) strongly favors the recovery of MscS back to the closed state. These may be common recovery mechanisms for the entire family of MscS-like channels with characteristic cage and kinked pore-lining helices, which evolved to function under turgor pressure exclusively in walled organisms. Perturbations (compromised state) of the cell wall lead to the conversion of turgor into membrane tension, which may intermittently lead to opening, but as a defensive action, it would rather drive the channel into the inactivated state.

## Acknowledgment

This work was supported by NIH R01-AI135015 to SS. The authors thank M. Britt, E. Moller, and J. Vanegas for suggestions and edits.

## References

Akitake, B., A. Anishkin, N. Liu, and S. Sukharev. 2007. Straightening and sequential buckling of the pore-lining helices define the gating cycle of MscS. Nat Struct Mol Biol. 14:1141–1149.

Akitake, B., A. Anishkin, and S. Sukharev. 2005. The “dashpot” mechanism of stretch-dependent gating in MscS. J.Gen.Physiol. 125:143–154.

Anishkin, A., B. Akitake, and S. Sukharev. 2008a. Characterization of the resting MscS: modeling and analysis of the closed bacterial mechanosensitive channel of small conductance. Biophys J. 94:1252–1266.

Anishkin, A., K. Kamaraju, and S. Sukharev. 2008b. Mechanosensitive channel MscS in the open state: modeling of the transition, explicit simulations, and experimental measurements of conductance. J Gen Physiol. 132:67–83.

Arnoldi, M., M. Fritz, E. Bauerlein, M. Radmacher, E. Sackmann, and A. Boulbitch. 2000. Bacterial turgor pressure can be measured by atomic force microscopy. Phys Rev E. 62:1034–1044.

Atilgan, E., V. Magidson, A. Khodjakov, and F. Chang. 2015. Morphogenesis of the Fission Yeast Cell through Cell Wall Expansion. Curr Biol. 25:2150–2157.

Belyy, V., A. Anishkin, K. Kamaraju, N. Liu, and S. Sukharev. 2010a. The tension-transmitting ‘clutch’ in the mechanosensitive channel MscS. Nat.Struct.Mol.Biol. 17:451–458.

Belyy, V., K. Kamaraju, B. Akitake, A. Anishkin, and S. Sukharev. 2010b. Adaptive behavior of bacterial mechanosensitive channels is coupled to membrane mechanics. J Gen Physiol. 135:641–652.

Boer, M., A. Anishkin, and S. Sukharev. 2011. Adaptive MscS gating in the osmotic permeability response in E. coli: the question of time. Biochemistry. 50:4087–4096.

Booth, I.R., and P. Blount. 2012. The MscS and MscL families of mechanosensitive channels act as microbial emergency release valves. Journal of bacteriology. 194:4802–4809.

Britt, M., E. Moller, J. Maramba, A. Anishkin, and S. Sukharev. 2024. MscS inactivation and recovery are slow voltage-dependent processes sensitive to interactions with lipids. Biophys J. 123:195–209.

Cayley, D.S., H.J. Guttman, and M.T. Record, Jr. 2000. Biophysical characterization of changes in amounts and activity of Escherichia coli cell and compartment water and turgor pressure in response to osmotic stress. Biophys J. 78:1748–1764.

Cox, C.D., N. Bavi, and B. Martinac. 2018. Bacterial Mechanosensors. Annu Rev Physiol. 80:71–93.

Flegler, V.J., A. Rasmussen, K. Borbil, L. Boten, H.A. Chen, H. Deinlein, J. Halang, K. Hellmanzik, J. Loffler, V. Schmidt, C. Makbul, C. Kraft, R. Hedrich, T. Rasmussen, and B. Bottcher. 2021. Mechanosensitive channel gating by delipidation. Proc Natl Acad Sci U S A. 118.

Hamilton, E.S., A.M. Schlegel, and E.S. Haswell. 2015. United in diversity: mechanosensitive ion channels in plants. Annu Rev Plant Biol. 66:113–137.

Hassoun, Y., J. Bartoli, A. Wahl, J.P. Viala, and E. Bouveret. 2021. Dual Regulation of Phosphatidylserine Decarboxylase Expression by Envelope Stress Responses. Front Mol Biosci. 8:665977.

Kamaraju, K., V. Belyy, I. Rowe, A. Anishkin, and S. Sukharev. 2011. The pathway and spatial scale for MscS inactivation. Journal of General Physiology. 138:49–57.

Koch, A.L., and M.F. Pinette. 1987. Nephelometric determination of turgor pressure in growing gramnegative bacteria. J.Bacteriol. 169:3654–3663.

Kung, C., B. Martinac, and S. Sukharev. 2010. Mechanosensitive channels in microbes. Annu.Rev.Microbiol. 64:313–329.

Lai, J.Y., Y.S. Poon, J.T. Kaiser, and D.C. Rees. 2013. Open and shut: crystal structures of the dodecylmaltoside solubilized mechanosensitive channel of small conductance from Escherichia coli and Helicobacter pylori at 4.4 A and 4.1 A resolutions. Protein Sci. 22:502–509.

Levina, N., S. Totemeyer, N.R. Stokes, P. Louis, M.A. Jones, and I.R. Booth. 1999. Protection of Escherichia coli cells against extreme turgor by activation of MscS and MscL mechanosensitive channels: identification of genes required for MscS activity. EMBO J. 18:1730–1737.

Martinac, B., and C. Kung. 2022. The force-from-lipid principle and its origin, a ‘what is true for E. coli is true for the elephant’ refrain. J Neurogenet. 36:44–54.

Moller, E., M. Britt, A. Schams, H. Cetuk, A. Anishkin, and S. Sukharev. 2023. Mechanosensitive channel MscS is critical for termination of the bacterial hypoosmotic permeability response. J Gen Physiol. 155.

Money, N.P., and R.J. Howard. 1996. Confirmation of a link between fungal pigmentation, turgor pressure, and pathogenicity using a new method of turgor measurement. Fungal Genet Biol. 20:217–227.

Park, Y.C., B. Reddy, N. Bavi, E. Perozo, and J.D. Faraldo-Gomez. 2023. State-specific morphological deformations of the lipid bilayer explain mechanosensitive gating of MscS ion channels. Elife. 12.

Pivetti, C.D., M.R. Yen, S. Miller, W. Busch, Y.H. Tseng, I.R. Booth, and M.H. Saier, Jr. 2003. Two families of mechanosensitive channel proteins. Microbiol.Mol.Biol.Rev. 67:66–85, table.

Pliotas, C., A.C. Dahl, T. Rasmussen, K.R. Mahendran, T.K. Smith, P. Marius, J. Gault, T. Banda, A. Rasmussen, S. Miller, C.V. Robinson, H. Bayley, M.S. Sansom, I.R. Booth, and J.H. Naismith. 2015. The role of lipids in mechanosensation. Nat Struct Mol Biol. 22:991–998.

Reddy, B., N. Bavi, A. Lu, Y. Park, and E. Perozo. 2019. Molecular basis of force-from-lipids gating in the mechanosensitive channel MscS. Elife. 8.

Rojas, E.R., K.C. Huang, and J.A. Theriot. 2017. Homeostatic Cell Growth Is Accomplished Mechanically through Membrane Tension Inhibition of Cell-Wall Synthesis. Cell Syst. 5:578–590 e576.

Rowe, I., A. Anishkin, K. Kamaraju, K. Yoshimura, and S. Sukharev. 2014. The cytoplasmic cage domain of the mechanosensitive channel MscS is a sensor of macromolecular crowding. J Gen Physiol. 143:543–557.

Sukharev, S. 2002. Purification of the small mechanosensitive channel of Escherichia coli (MscS): the subunit structure, conduction, and gating characteristics in liposomes. Biophys.J. 83:290–298.

Tyree, M.T., and H.T. Hammel. 1972. Measurement of Turgor Pressure and Water Relations of Plants by Pressure-Bomb Technique. J Exp Bot. 23:267-&.

Whatmore, A.M., and R.H. Reed. 1990. Determination of turgor pressure in Bacillus subtilis: a possible role for K+ in turgor regulation. J Gen Microbiol. 136:2521–2526.

Wong, F., S. Wilson, R. Helbig, S. Hegde, O. Aftenieva, H. Zheng, C. Liu, T. Pilizota, E.C. Garner, A. Amir, and L.D. Renner. 2021. Understanding Beta-Lactam-Induced Lysis at the Single-Cell Level. Front Microbiol. 12:712007.

Zhang, Y., C. Daday, R.X. Gu, C.D. Cox, B. Martinac, B.L. de Groot, and T. Walz. 2021. Visualization of the mechanosensitive ion channel MscS under membrane tension. Nature. 590:509–514.

